# Cotranslational Protein Folding Through Non-Native Structural Intermediates

**DOI:** 10.1101/2025.04.09.648002

**Authors:** Siyu Wang, Amir Bitran, Ekaterina Samatova, Eugene I. Shakhnovich, Marina V. Rodnina

**Affiliations:** Department of Physical Biochemistry, Max Planck Institute for Multidisciplinary Sciences; Göttingen, 37077, Germany; Department of Chemistry and Chemical Biology, Harvard University; 12 Oxford Street Cambridge, Massachusetts 02138; Department of Molecular and Cellular Biology, University of California Berkeley, Berkeley CA 94720

## Abstract

Cotranslational protein folding follows a distinct pathway shaped by the vectorial emergence of the peptide and spatial constraints of the ribosome exit tunnel. Variations in translation rhythm can cause misfolding linked to disease; however, predicting cotranslational folding pathways remains challenging. Here we computationally predict and experimentally validate a vectorial hierarchy of folding resolved at the atomistic level, where early intermediates are stabilized through non-native hydrophobic interactions before rearranging into the native-like fold. Disrupting these interactions destabilizes intermediates and impairs folding. The chaperone Trigger Factor alters the cotranslational folding pathway by keeping the nascent peptide dynamic until the full domain emerges. Our results highlight an unexpected role of surface-exposed residues in protein folding on the ribosome and provide tools to improve folding prediction and protein design.

---

In all cells, proteins begin to fold into their functional structures during synthesis on the ribosome (*1–4*). Unlike folding in solution, where the full sequence is available, cotranslational folding occurs vectorially as the nascent protein emerges from the ribosome exit tunnel, allowing early folding before synthesis is complete (*5–12*). The ribosome orchestrates this process by imposing spatial constraints, interacting with the nascent peptide, and modulating the structural dynamics of the emerging domains, which collectively shape the folding pathway. Interactions with ribosomal components, including rRNA and proteins lining the exit tunnel, influence the folding trajectory by stabilizing specific conformations and regulating structural transitions (*6, 12–20*). Upon emerging from the ribosome and release into the cytosol, some proteins can undergo further folding/refolding cycles with or without the help of molecular chaperones, while others are not refoldable (*21–24*). Disruptions of cotranslational folding caused by variations in local translation rates or mutations can lead to misfolding and disease (*25–31*), highlighting the importance of this process for protein homeostasis. While many proteins start folding inside the ribosome exit tunnel, the guiding principles of cotranslational folding remain unclear, as intermediates are transient, and ribosomal interactions complicate structural determination. Here, we combine computational and experimental approaches to reconstruct the cotranslational folding pathway using the N-terminal domain of HemK as a model protein. HemK NTD is a five-helix bundle (Fig. 1A) that folds rapidly in a two-state concerted transition in solution, whereas on the ribosome, folding is sequential, involving α-helix formation and stepwise compaction (*9, 20, 32*). We gain insights into atomistic structure and dynamics of cotranslational intermediates, revealing significant non-native interactions, characterize their formation and conversion into final native structure, and demonstrate the role of the chaperone trigger factor (TF) in resolving non-native interaction during cotranslational folding.

**Fig. 1.**
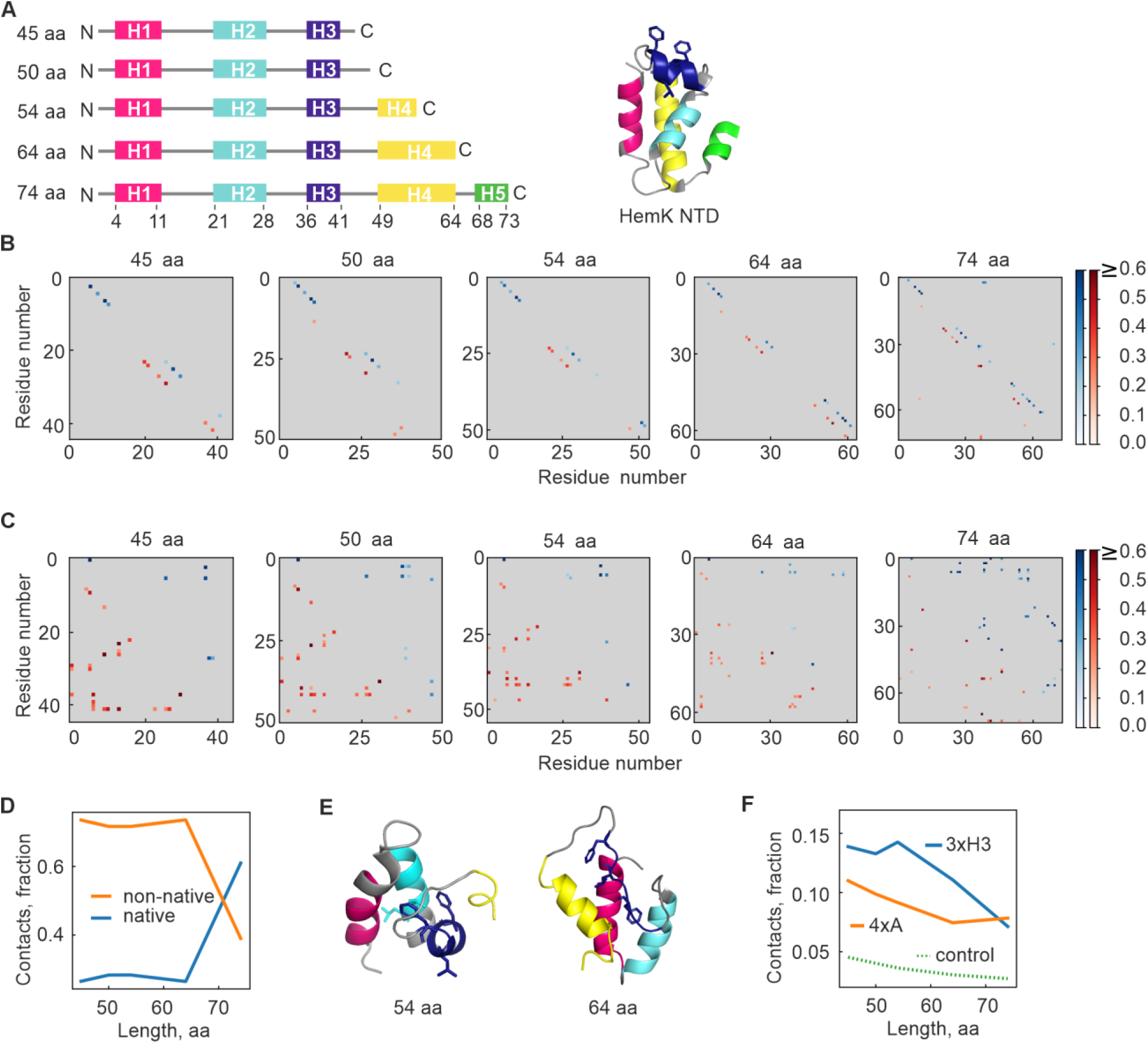
Simulations predict an early non-native folding intermediate of HemK NTD stabilized by hydrophobic interactions. (**A**) Schematic representation of simulated constructs, with helices labeled H1 to H5. Full-length HemK NTD (aa 1-73) forms a stable, autonomously folding domain of five α-helices. (**B**) Average contact maps at various chain lengths for α-carbon interactions. Blue dots (top-right diagonal) represent native helical contacts between residues *i* and *i*+3, while red dots (bottom-left diagonal) indicate slightly register-shifted, non-native helical contacts. Native contacts (blue, upper diagonals) and non-native contacts (red, lower diagonals) are scaled by probability (color bar) with contacts greater than 0.2 shown. (**C**) Same as (B) but for side-chain contacts (≤ 5 Å distance between side chain centers) at different protein lengths. (**D**) Fraction of native vs. non-native side-chain contacts at each protein length. (**E**) Sample simulation snapshots of non-native intermediates at 54 aa and 64 aa chain length. 3xH3 residues (F38, L40 or F42) are shown in as dark blue sticks. (**F**) Fraction of sidechain contacts at each length involving 3xH3 (blue) or 4xA (L27, L28, L55 or L58, orange). Values are normalized by the number of residues in each group. For control, residue numbers were randomly shuffled, and 3xH3 contact frequencies recalculated.

## RESULTS

### Computational Predictions of Folding Hierarchy

To identify key residues guiding this process, we performed atomistic simulations using MCPU/DBFOLD, a Monte Carlo simulation pipeline that predicts protein folding pathways while accounting for non-native interactions (*33*). We first validate that the wild-type (WT) full-length HemK NTD is well-folded at low temperatures and undergoes a cooperative two-state thermal denaturation in our simulation, consistent with experiments (*9*) (fig. S1A). Although the computational platform does not explicitly model the ribosome, these simulations recapitulate intrinsic properties of cotranslational folding, such as the sequential emergence of the peptide and its propensity to form secondary and tertiary structures. They also provide insights into the thermodynamic stability and dimensions of folding intermediates, which determine whether structural elements can form inside the narrow ribosome exit tunnel. To mimic the vectorial nature of cotranslational folding, we ran equilibrium simulations of the domain truncated at various C-terminal positions (Fig. 1A). At intermediate lengths (45–64 amino acids (aa)), the protein readily forms native-like α-helices (Fig. 1B), but adopts persistent non-native tertiary structures (∼70% non-native contacts, Fig. 1B,C), which are favored over native-like structures by 2-3 kBT (fig. S1B,C) Native contacts become more important only at full-length (74 aa) (Fig. 1D), with a stability gain of ∼4 kBT relative to the unfolded state (fig. S1C).

Next, we analyzed molecular interactions stabilizing these intermediates. Three hydrophobic residues in helix 3 (Phe38, Leu40, Phe42, hereafter denoted as 3xH3), which are typically solvent-exposed in the native state, are frequently buried in non-native intermediates (Fig. 1E). At short chain lengths, 3xH3 residues account for ∼15% of side-chain contacts, exceeding random expectations (Fig. 1F). In contrast, native hydrophobic core residues (Leu27, 28, 55, 58, previously denoted as 4xA (*9*)) account for only ∼10% of interactions at shorter lengths, but become dominant at later stages. At full length, 4xA residues dominate, stabilizing the native state, although non-native 3xH3 contacts are still visible, likely due to structural fluctuations that transiently bury these hydrophobics within the core, despite their solvent-exposed nature in the crystal structure. To assess their role, we reran simulations with mutants in which either 3xH3 or 4xA residues were replaced with Ala. Surprisingly, at full length, both mutations destabilized the native state—despite 3xH3 residues being expected to become surface-exposed—although the destabilizing effect was more pronounced for the hydrophobic core mutation, 4xA (fig S1C-E), consistent with the observed hierarchy of contact formation (Fig. 1F). Moreover, 3xH3, but not the 4xA, affected the stability of folding intermediates prior to completion of synthesis (fig S1B-D). These results suggest a vectorial hierarchy in cotranslational intermediates, with 3xH3 residues initially forming a non-native core that rearranges toward native-like, albeit destabilized, domain as translation progresses.

### Non-Native Interactions in Cotranslational Folding

We then experimentally tested the simulation predictions using Fluorescence Correlation Spectroscopy (FCS) combined with fluorescence quenching via photo-induced electron transfer (PET). We used a fluorescence label (Atto655) at Met1, which is quenched upon contacting Trp6 in the nascent chain and becomes unquenched as it moves away, reflecting protein structural dynamics in the µs to ms time range (*34, 35*) (Fig. 2A). To test its sensitivity to hierarchical folding, we calculated the root-mean-square fluctuation (RMSF) between residues 1 and 6 for the WT and 3xH3 and 4xA mutants. Simulations predict that 3xH3 mutations increase dynamics when the nascent peptide is short while their effect on dynamics diminishes at full length, whereas 4xA mutations affect dynamics at full length (Fig. 2B), consistent with their role in native folding (*9*).

**Fig. 2.**
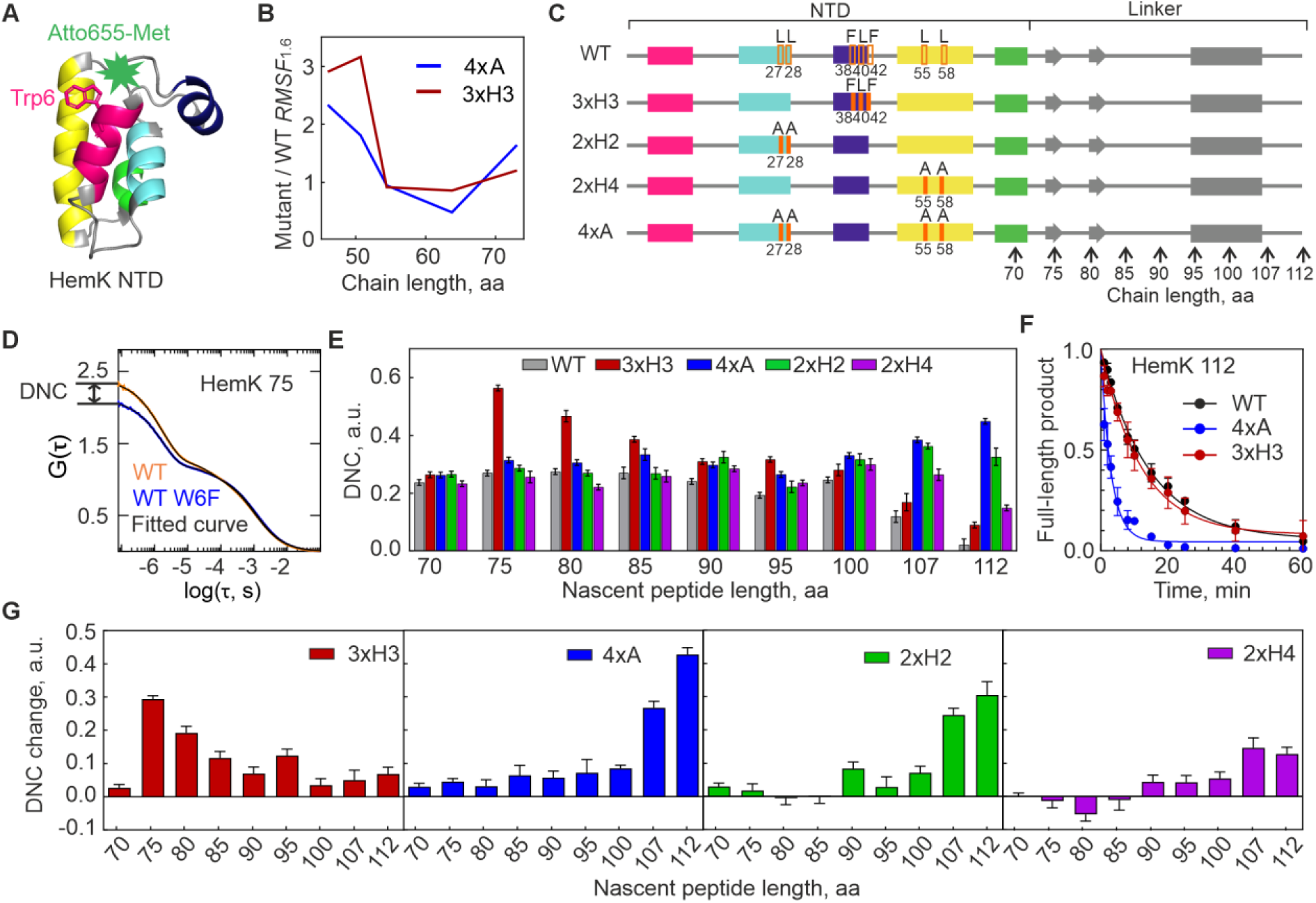
HemK NTD forms non-native intermediates during translation. (**A**) Structure of HemK NTD (PDB 1T43) with the N-terminal fluorophore highlighted in green and Trp6 shown in magenta (stick representation). (**B**) Predicted RMSF in the distance between residues 1 and 6 for 3xH3 and 4xA mutants relative to WT at equivalent nascent chain lengths. (**C**) Schematic of constructs for PET experiments, including the NTD (aa 1-72) and the linker (gray, aa 73-112) connecting the emerging domain to the peptidyl transferase center of the ribosome. As linker, native HemK sequence was used, which entails potential secondary structure elements such as two short β-strands and an α-helix as indicated. Vertical arrows indicate the aa length of nascent-chain constructs used in the experiments. Orange rectangles indicate mutation sites. (**D**) Example of PET-FCS data with fitted curves (black). Dynamicity of the nascent chain (DNC), a qualitative parameter used to assess the fraction of molecules undergoing conformational fluctuations, was determined from the G(0) difference between W6 and W6F ribosome-nascent chain constructs. (**E**) DNC values for all tested constructs, determined from the G(0) differences as described in (**D**); error bars represent SEM. (**F**) Protease digestion of HemK112 nascent chains. Full-length peptide intensity at time zero was used for normalization. Data represent mean values with SD of three biological replicates. Smooth lines represent single-exponential fits used to calculate the decay constant (τ). τ = 14.4 ± 3.7 min (WT), 12.0 ± 1.2 min (3xH3), and 3.0 ±0.3 min (4xA) (mean ± SD). (**G**) DNC changes for the 3xH3, 4xA, 2xH2, and 2xH4 mutants relative to the WT. Error bars are SEM.

To measure the effects of mutations, we prepared ribosome-nascent chain complexes (RNCs) with WT, 3xH3, and 4xA constructs (Fig. 2C and fig. S2; Methods). Additionally, we designed double Ala mutants in helix H2 (Leu27Ala and Leu28Ala, denoted as 2xH2) and helix H4 (Leu55Ala and Leu58Ala, denoted as 2xH4) to assess individual contributions to folding. Since compact tertiary structures comparable to those observed in computer simulations form only after a portion of the peptide exits the ribosome, we tested constructs of different lengths, starting from 70 aa, which is completely buried inside the tunnel, up to 112 aa, which is known to expose the whole HemK NTD outside the ribosome (*9*) (Fig. 2C). To estimate the contribution of the ribosome-induced quenching effects, we prepared control constructs (W6F) where Trp6 was replaced by a non-quenching Phe. PET-FCS autocorrelation functions (ACF, denoted as G(τ) in Fig. 2D) showed a millisecond diffusion time identical for all RNCs (*20*) and a biphasic PET signal spanning the µs timescale. The higher signal for W6 compared to W6F peptides reflects nascent chain dynamics (fig. S3A,B). The Y-intercepts (G(0)) of ACF curves, representing quenching event frequency (*36*), enabled a simple estimation of the dynamicity of nascent chain (DNC) values from the difference between W6 and W6F ACFs (Fig. 2E; Methods). Higher DNC values indicate dynamic nascent chains, while stably folded molecules have values close to zero (*20*).

The WT nascent chains remain dynamic from 70 to 105 aa, gradually compacting beyond 107 aa as the NTD emerges from the exit tunnel and folds, in agreement with previous findings (*20*) (Fig. 2E,F and fig. S3C). The 3xH3 mutant exhibits WT-like dynamics at 70 aa but becomes significantly more dynamic at 75–100 aa, indicating destabilization of intermediates. At longer lengths, both WT and 3xH3 reach compact states, although 3xH3 remains somewhat more dynamic (Fig. 2E,G, consistent with the prediction of fig. S1C,D). Mutations in 4xA residues have distinctly different effects. The 2xH2, 2xH4, and 4xA mutations have little impact on early folding, with only mild destabilization between 90–100 aa. This supports the idea that hydrophobic core residues do not contribute to early folding intermediates (*20*). However, destabilization becomes significant beyond 107 aa, coinciding with the emergence of the domain from the ribosome tunnel. The effect is most pronounced for the mutations in helix 2, likely disrupting early H1-H2 interactions (*32*). Limited proteolysis assays indicate that the full-length 3xH3 mutant is compact (Fig. 2F), whereas 4xA is strongly destabilized (Fig. 2G and (*9*)). These findings indicate that 3xH3 residues stabilize early non-native intermediates, while 4xA residues are crucial for native state folding of the full-length domain, in agreement with computational predictions.

### Impact of Non-Native Interactions on Nascent Chain Compaction

On the ribosome, 3xH3 mutation disrupts folding into a non-native intermediate when the nascent chain reaches ∼75 aa. This non-native intermediate includes a ∼50 aa compact folding unit and a ∼ 25 aa linker (*9, 32*), comprising an intermediate with helix 3 docking onto a unit formed by native contacts between helices 1 and 2 (Fig. 1), which coincides with a major force-generating event during cotranslational folding of HemK NTD (*20*). To characterize this intermediate, we analyzed domain-wide dynamics at the atomic level by calculating the RMSF of pairwise distances between all amino acids relative to their average distance (Fig. 3). Although the structures of HemK (full-length or intermediates) are much more stable in solution than on the ribosome (*20*), such comparisons provide insights into the relative effects of mutations. In WT HemK NTD, RMSF values at intermediate peptide lengths largely match those of the full-length protein, with only a few localized fluctuations. In contrast, the 3xH3 mutant at lengths corresponding to the ∼50 aa folding unit (45–55 aa) shows significantly higher RMSF (Fig. 3A,B), particularly in the linker between helices 2 and 3 (residues 32–37) and at the C-terminus, suggesting that non-native interactions play a role in stabilizing folding intermediates as they emerge from the ribosome.

**Fig. 3.**
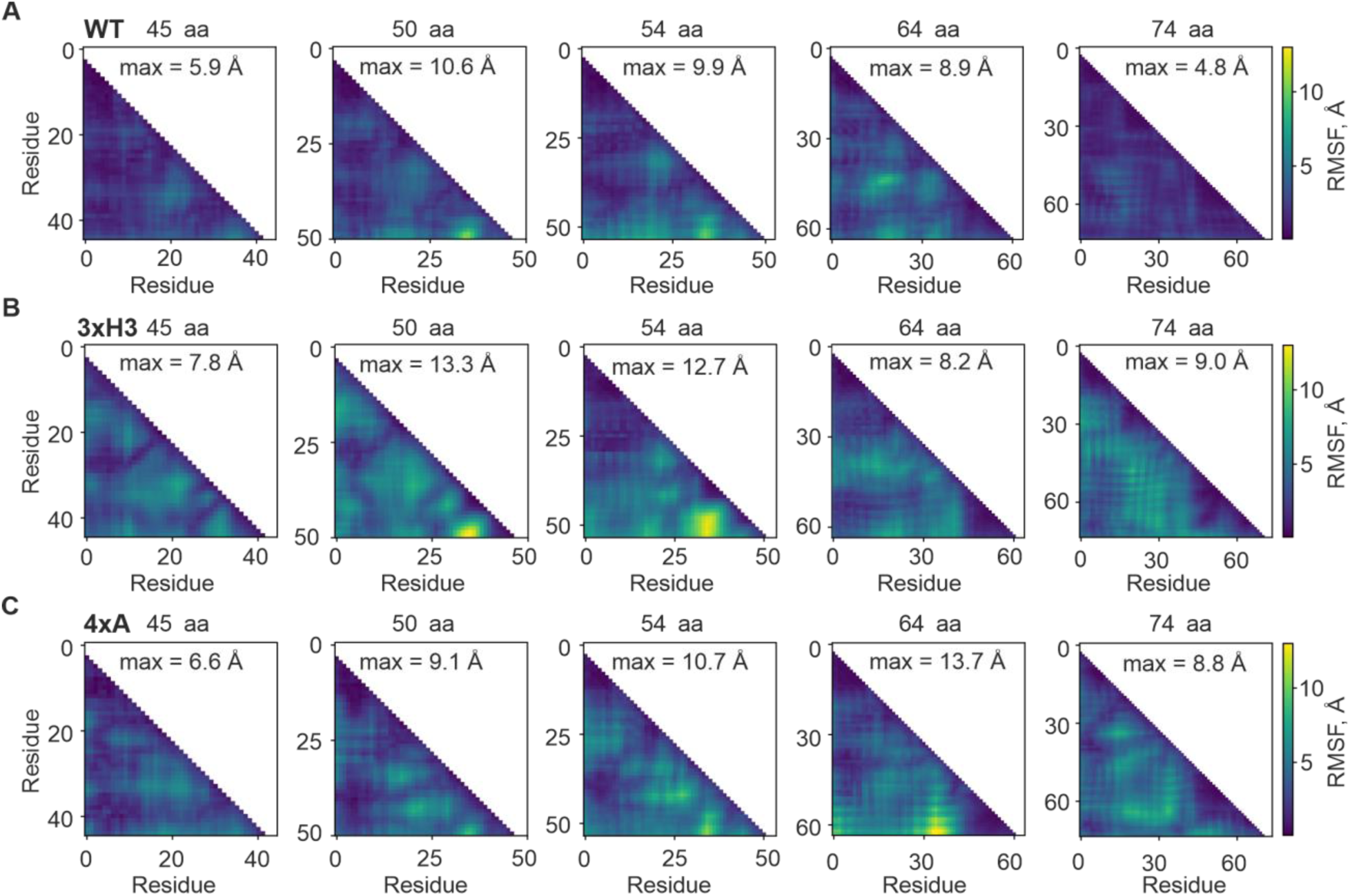
Disrupting non-native interactions destabilizes full-length domain folding. Heatmaps show RMSF, defined as the root mean squared fluctuation in the distances between residue pairs relative to their average distances, at different nascent chain lengths. RMSF values are color-coded according to the color bars to the right of each row. The maximum RMSF value in each heatmap is indicated. (**A**) WT HemK NTD. (**B**) 3xH3. (**C**) 4xA.

These findings are further supported by calculations of the radius of gyration (Rg) distribution for each length (fig. S4), representing the degree of domain compactness. WT HemK NTD remains consistently compact, with a mean Rg at 45–50 aa differing by no more than 1Å from the native structure and exhibiting a narrow Rg distribution (<2Å). This indicates that WT intermediates are compact despite the non-native interactions that stabilize them. The 3xH3 mutant, however, adopts more expanded states with a broad Rg distribution at 45–50 aa, reinforcing the idea that non-native interactions help form a compact intermediate suitable for helix 3 docking inside the exit tunnel. In contrast, the 4xA mutation has minimal impact on RMSF (Fig. 3C) and Rg distribution (fig. S4A-D) at short nascent peptide lengths, but its effect becomes more pronounced at 64 and 74 aa, where the protein transitions into a native-like state stabilized by 4xA residues.

### Real-Time Analysis of Folding

Although the mutated residues in the 3xH3 mutant are largely solvent-exposed in the native structure, the full-length 3xH3 protein appears more dynamic than the WT in both computer simulations (Fig. 3B, fig. S1D and S4E) and PET-FCS measurements (Fig. 2G). Furthermore, simulations indicate that the full-length 3xH3 mutant exists as a dynamic ensemble of conformations, including favorable non-native interactions (Fig. 3B, figs. S1, S4). On the other hand, protease protection experiments suggest a compact fold comparable to the WT (Fig. 2F and fig. S3C). This apparent contradiction prompted us to investigate the folding pathway using an orthogonal approach. While FCS captures ms-scale dynamics and protease protection assays reflect min-scale stability, we bridged these timescales by measuring real-time PET efficiency changes using a stopped-flow apparatus. We start translation by rapid mixing of translation elongation factors and aminoacyl-tRNAs with the ribosome in complex with an mRNA and BodipyFL-Met-tRNA^fMet^ and monitor changes in Bodipy fluorescence as the nascent peptide is synthesized. As for PET-FSC experiments, we perform these experiments with W6 and W6F peptides to distinguish between the contributions of the ribosome and nascent chain folding ((*20*) and Methods). During WT HemK NTD synthesis, PET efficiency initially increases, suggesting the formation of an intermediate where the fluorescent reporter is near Trp6, then decreases (Fig. 4A). The decrease coincides with full-length protein synthesis (fig. S5A,B), suggesting the transition to a native-like structure (*20*). As expected, the 4xA mutant does not undergo this final rearrangement (Fig. 4A). The full length 3xH3 mutant is also trapped in a non-native (Fig. 4A) yet compact (Fig. 2F) conformation, consistent with simulation results (Fig. 3B, figs. S1, S4). These results together with computer simulations (Fig. 3) demonstrate that disrupting non-native hydrophobic interactions which guide hierarchical folding can lead to a compact but partially misfolded protein.

**Fig. 4.**
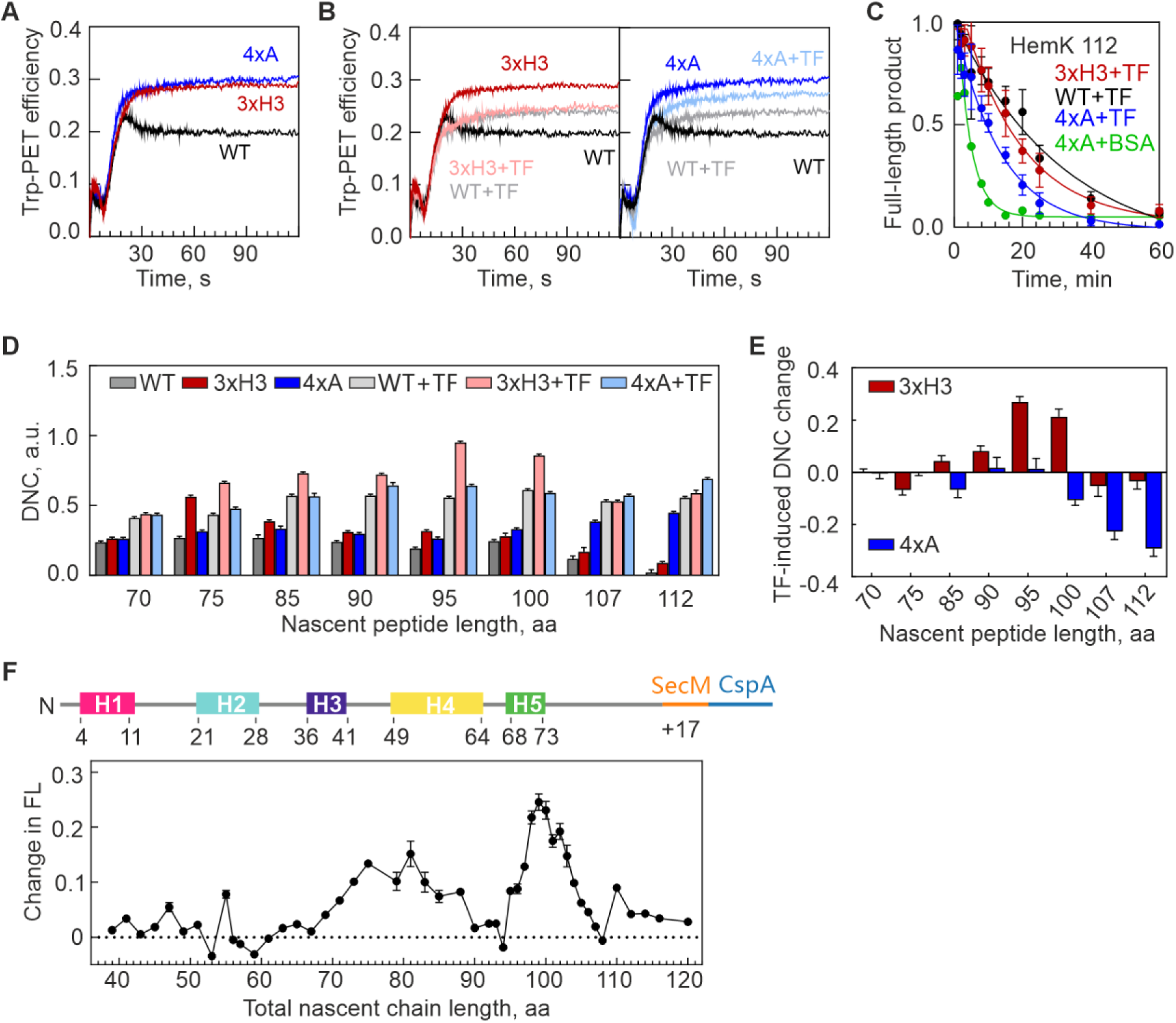
TF alleviates misfolding by destabilizing intermediates and the final domain fold. (**A**). Stopped-flow time course of HemK NTD cotranslational compaction monitored by PET between Atto655-Met1 and Trp6 in the nascent chain. Time zero corresponds to translation start upon mixing the 70S–mRNA–Atto655-Met-tRNA^fMet^ complex with EF-Tu, EF-G and aminoacyl-tRNAs. (**B**) TF effect on cotranslational folding of WT, 3xH3 (left panel) and 4xA (right panel). Note that the traces for WT+TF (light gray) and 3xH3 (light red) are almost identical. (**C**) Time courses of protease digestion in the presence of TF. Data represent mean ± SD of three biological replicates. τ = 31.7± 6.0 min (WT+TF), 22.4 ± 3.1 min (3xH3+TF), 13.5 ± 1.5 min (4xA+TF), and 3.7 ±0.7 min (4xA+BSA). (**D**) TF effect on DNC monitored by PET-FSC as in Fig. 2. (**E**) Relative TF-induced DNC change for 3xH3 and 4xA compared to the WT+TF nascent chain of the same length. Plotted is the dynamicity change upon TF addition of the mutant vs. WT. (**F**) TF-induced change in the FPA profile of WT nascent peptide. The total nascent chain length includes the HemK nascent peptide and the SecM stalling sequence. Data represent mean ± SD of three biological replicates.

### Role of Trigger Factor

In cells, molecular chaperones prevent protein misfolding. This prompted us to ask whether chaperones can also correct cotranslational misfolding caused by the alterations in folding pathway. Trigger Factor (TF), a co-translational chaperone that binds at the ribosome tunnel exit, interacts directly with nascent peptides (*37*). In real-time translation experiments, TF prevents relaxation of the WT HemK domain (Fig. 4B), with effects visible after 10-14 s (fig. S5C) corresponding to the synthesis of the first 70-90 aa (fig. S5B). This suggests that TF engages early, influencing folding by binding to a crucial intermediate that forms during the synthesis. When TF is added during 3xH3 mutant translation, PET efficiency becomes identical to that of the WT (Fig. 4B), eliminating the folding differences between the WT and 3xH3 variant. Notably, the overall domain structure remains compact (Fig. 4C). These data indicate that by binding to an early intermediate, TF does not block the overall domain compaction, but delays final folding, which might avoid formation of the compact misfolded forms arising when the 3xH3 interactions cannot form. In contrast, while TF affects structure of the 4xA mutant (Fig. 4B, C), its final PET efficiency in 4xA remains higher than in WT, suggesting that TF cannot eliminate the folding defect that arises when the hydrophobic core is disrupted.

To examine how TF affects protein fluctuations on a µs timescale, we performed PET-FCS experiments. TF increases dynamicity in both WT and mutant proteins at all peptide lengths (Fig. 4D, fig. S5D,E), supporting the idea that TF destabilizes nascent peptide folding on the ribosome. However, the magnitude of the effect varies with the nascent chain: TF increases fluctuations in 3xH3 more than in the WT at 95-100 aa (Fig. 4E). For 4xA, the effect is similar to that of the wild-type (WT) up to a chain length of 100 amino acids. However, at longer chain lengths, the effect is reduced compared to WT, possibly due to stabilization by bound TF of a domain with a defective hydrophobic core (Fig. 4C, Fig. S5F).

As stopped-flow kinetics indicated early TF recruitment to the nascent chain, while PET-FCS suggested its main effect occurs at longer chain lengths, we examined TF’s influence on folding using an orthogonal approach. We performed high-resolution force profile analysis (FPA) (*5*) using constructs encoding HemK NTD peptides of varying lengths (8 to 103 amino acids), each followed by 17 codons of the SecM arrest peptide and a 20-aa reporter (*20*). In the absence of tension, translation stalls at the SecM arrest motif. When mechanical force is generated, translational arrest is relieved, allowing synthesis to continue. Without TF, the FPA profile reveals distinct peaks corresponding to the sequential folding of the NTD through stepwise helix formation and docking (fig S5G, S6) (*20*). TF increases tension, particularly at nascent chain lengths of 72– 87 aa and 95–103 aa (Fig. 4F, fig. S5G and fig. S6). Once the peptide reaches 112 amino acids, both profiles converge at a baseline level, indicating that the emerged NTD does not exert tension on the ribosome regardless of TF’s presence. The increased tension at 72–87 aa coincides with TF’s early recruitment observed in stopped-flow experiments, while the peak at 95–103 aa overlaps with the length at which the maximum destabilizing effect of TF on folding is observed. This additional tension reflects mechanical force preventing the final compaction of the full-length nascent chain as it emerges from the ribosome.

## DISCUSSION

Our simulations predict cotranslational folding intermediates at atomistic level of detail. Experimental data confirm the presence of these intermediates, revealing that the cotranslational folding of HemK NTD involves well-defined structures stabilized by specific non-native contacts as the nascent peptide emerges from the ribosome. The temporal hierarchy of these intermediates defines the folding pathway, guiding the nascent chain towards its final native conformation. These non-native intermediates may help compact the nascent chain within the narrow ribosomal tunnel, facilitating proper folding. Indeed, our Rg analysis suggests that without these interactions, nascent chains would be less compact, which could hinder folding and increase the risk of misfolding. Mutations that disrupt critical non-native interactions not only alter the folding pathway but also result in a different ensemble of conformations at full length. While the equilibrium in this ensemble is shifted towards a compact state, as shown by protease protection experiments, the local interactions remain dynamic, suggesting local misfolding. TF engages with nascent chains early during translation, at a stage when hydrophobic interactions begin to form a non-native compact intermediate, and modulate the nascent chain dynamics of the growing peptide. TF destabilizes both native-like and potentially misfolded structures, thereby delaying final folding (*38*). The mechanical force generated by interactions between TF, the nascent peptide, and the ribosome destabilizes prematurely folded structures, ensuring that the peptide remains flexible and capable of adopting its correct native conformation once the complete domain sequence emerges from the ribosome and the nascent chained is released from the ribosome and TF. We show that TF can mitigate cotranslational misfolding even in proteins that are not obligate TF clients, such as HemK, by delaying their final compaction into a stable fold. Once the nascent chain is released from both the ribosome and TF, these proteins have a chance to attain their stable correct structure. Misfolding due to altered folding pathways may become critical under stress conditions, when chaperone capacity is overwhelmed. This is particularly important for proteins that are inherently unable to refold in solution (*22*), as cotranslational TF-mediated control of their final packing may represent the last opportunity for refolding before potentially misfolded protein is released into the cytoplasm.

While the native fold is encoded in the sequence, evolutionary adaptation of cotranslational folding likely involves residues that do not contribute to the final structure but are essential for correctly folding intermediates, reflecting the vectorial nature of protein synthesis and the interactions of the nascent chains with the ribosome exit tunnel. Replacing such seemingly neutral surface residues can disrupt intermediates and lead to misfolding. This insight is crucial for understanding how translation rhythm influences protein folding and for designing novel protein structures. For example, this may explain why variations in translation rates at codons encoding amino acids that do not directly participate in folding can cause disease and why mutations of residues that are exposed on the protein surface in the native structure can still be detrimental to proper folding. Our results also demonstrate that simulations of vectorially truncated N-terminal peptides folding in solution can predict intermediate structures as peptides exit the ribosome. This approach enables the construction of cotranslational folding profiles for any protein, including large, misfolding-prone ones, and can model interactions with chaperones, biogenesis factors, or subunits. These findings advance our understanding of protein folding on the ribosome and open new avenues for future research.

## Acknowledgments

We thank Olaf Geintzer, Vanessa Herold, Franziska Hummel, Sandra Kappler, Christina Kothe, Anna Pfeifer, and Michael Zimmermann for expert technical assistance. The authors used chatgpt4o to improve the readability of the manuscript. After using chatgpt4o, the authors reviewed and edited the content as needed and take full responsibility for the content of the manuscript.

## Author contributions

Conceptualization: AB, ES, EIS, MVR

Methodology: SW, AB, ES

Investigation: SW, AB

Visualization: SW, AB

Funding acquisition: EIS, MVR

Project administration: ES, EIS, MVR

Supervision: ES, EIS, MVR

Writing – original draft: SW, AB, ES, EIS, MVR

Writing – review & editing: SW, AB, ES, EIS, MVR

## Data and materials availability

All data are available in the main text or the supplementary materials.

## Supplementary Materials

Materials and Methods

Figs. S1 to S6

Tables S1 to S2

References (*39–46*)

## MATERIALS AND METHODS

### Computational Methods

#### All-atom Monte-Carlo Simulations

Simulations of all HemK constructs were carried out using the MCPU, an all-atom Monte-Carlo (MC) simulation algorithm that uses a knowledge-based potential function to compute interaction energies between all residues’ pairs, account for both native and nonnative interactions, as described previously (*33*). We prepared starting structures for simulations from the PDB file 1T43, with Arg at position 34 (Arg34) replaced with Lys to match the construct used in experiments. Residues 75 onwards were deleted to produce a 74 amino acids (aa) HemK NTD fragment containing all five helices; this construct is hereafter referred to as full-length HemK NTD. We then generated additional truncations from the C-terminus to produce variable-length N-terminal fragments as described in the text. This process was repeated to generate full-length and truncated HemK variants with 3xH3 (F38, L40 or F42) and 4xA (L27, 28, 55 or 58) replacements with Ala. The sequences for the respective constructs were used to generate sequence-specific parameters for local conformational energy calculations as described (*39*). Each starting structure was equilibrated in the MCPU potential function using the low-temperature replica-exchange equilibration protocol (*33, 40*) with a distance cutoff of 8.5Å used to compute α-carbon contacts for umbrella biasing. A low simulation temperature of 0.1 (in units of simulation energy) was used to relax the structure, and following 20 million MC steps of equilibration, the lowest energy structure amongst all replicas was identified and used as the starting structure for subsequent production simulations of the respective construct.

Production runs were then carried out for 200 million MC steps at a range of temperatures from T=0.4 to T=1.0 in increments of 0.025, and umbrella biasing setpoints ranging from 0 to the number of α-carbon contacts in the equilibrated structure, in increments of 10 contacts, producing a grid of simulations in setpoint and temperature space. Replica exchanges were periodically attempted between neighboring simulations as in (*33*) to help improve convergence. Following completion of the 200 million MC step simulation, the second 100 million steps were used to calculate equilibrium properties. This was justified by the observation that the sliding-window averaged energy as a function of simulation step generally ceased changing appreciably after 100 million MC steps, indicating convergence.

### Computation of thermodynamic properties from all-atom simulations

To analyze converged simulations for each construct, we began by computing the average number of native contacts, defined as contacts that present in the equilibrated native structure, as a function of simulation temperature using MBAR (*41*), a statistically-optimal approach for estimating thermodynamic properties from biased simulations. This method can be used to calculate thermal melting curves like those shown in fig. S1A. From this melting curve, we identified the melting temperature for WT full-length NTD as TM=0.525—the temperature at which the average number of native contacts is closest to the half-maximal value. We then constructed more detailed folding landscapes using the substructure approach detailed in (*33*).

In this method, clusters of cooperatively-forming native contacts are identified from the contact map—in the case of HemK, we used an 8.5 Å distance cutoff for contacts excluding residues less than eight apart in primary sequence, and we clustered together contacts separated by no more than four Manhattan distance units on the contact map, provided the resulting clusters were at least seven residues in size. The resulting clusters, known as *substructures* are labeled A-E in fig. S1E. Each simulation snapshot was then assigned to a *topological configuration,* which is a label that identifies the substructures that are formed or broken in that particular snapshot. We then computed a potential of mean force (PMF) as a function of topological configuration using MBAR as in fig. S1C-E.

As an alternative approach to characterize native vs nonnative folding propensities, we generated average contact maps at a fixed temperature of T=0.45 (or T=0.86 TM with TM=0.525) as shown in Fig. 1. Two types of contact maps were generated. In the first, we computed centroid positions for each side chain in every snapshot. We then generated a side-chain contact map for each snapshot by identifying all pairs of side chains for which the distance between the respective centroids is less than 5Å, provided they are at least four residues apart in primary sequence. We then computed a thermally averaged contact map using MBAR. Alternatively, the same calculation was performed using α-carbons with the same distance cutoff. From side chain contact maps, we identified native contacts as those within one Manhattan distance unit of contacts present in the equilibrated native structure—this distance tolerance allows slightly register-shifted contacts to still be counted as native-like. This allows us to calculate the fraction of contacts in each snapshot that are native or nonnative. We also used these contact maps to count the total number of contacts involving 3xH3 or 4xA residues, and these values were normalized by the number of residues in each group. As a control, this calculation was repeated for 3xH3 residues in contact maps where residue indices were randomly reshuffled. All these quantities were thermally averaged across snapshots using the MBAR algorithm.

For each snapshot from our converged simulation at temperature T=0.45, we computed the radius of gyration as follows:

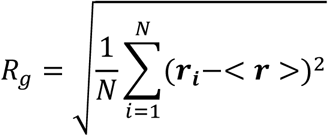

where i indicates residues, N is the total number of residues in the construct, ***r_i_*** is the position of the i^th^ α-carbon, and <**r**> is the average position of all α-carbons in the structure. This quantity was computed separately for each snapshot, then MBAR was used to obtain a probability distribution P(*R*_*g*_) and average <*R*_*g*_> over snapshots. These quantities are shown in fig. S4. As a control, we calculated the average Rg that would be expected for a perfectly disordered random coil as

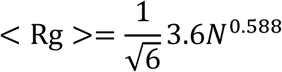

where 3.6 is the approximate contour length of an individual amino acid (in angstroms), N is the number of residues, and 0.588 is the scaling exponent for the end-to-end distance of a self-avoiding random walk in 3D (*42*). These values are shown in fig. S4.

Finally, as a metric for dynamicity of a given construct, we used MBAR to calculate, for each residue pair, the distribution of distances between the respective alpha carbons. From these distributions we calculated the mean squared fluctuation relative to the average distance, and taking the square root of this quantity gives the room-mean squared fluctuation or RMSF. These RMSF values are shown for each residue pair in Fig. 3. The pairwise value corresponding to residues 1 and 6 was used to generate Fig. 2B.

## Biochemical Methods

### Buffers and reagents

Biochemical experiments were carried out in HiFi buffer (50 mM HEPES–HCl, pH 7.5, 70 mM NH4Cl, 30 mM KCl and 3.5 mM MgCl2, 8 mM putrescine and 0.5 mM spermidine) or HAKM7 buffer (50 mM HEPES–HCl, pH 7.5, 70 mM NH4Cl, 30 mM KCl and 7 mM MgCl2) (*43*). 70S ribosomes and other translational components, including initiation factors (IF1, IF2, IF3), initiator tRNA (tRNA^fMet^), total aminoacyl-tRNA (aa-tRNA), EF-G, EF-Tu, and EF-Ts, were prepared according to published protocols (*44, 45*).

### Plasmid construction

The coding sequence of HemK NTD was cloned into pEX-A128 vector, which carries an ampicillin resistance cassette (Eurofins Genomics, Ebersberg, Germany). For PET-FCS measurements, the native Trp at position 6 (W6) was used, while Trp78 and Tyr3 were mutated to Phe in all WT and mutant constructs to avoid PET interactions with the N-terminal fluorescent reporter.

For the Force Profile Assay (FPA), a series of constructs containing different length of WT HemK (aa 1-101) were designed, each followed by the 17-aa SecM arrest peptide (*46*) and 20-aa cold shock protein A (CspA) as reporter (UniProt ID: P0A9X9) were prepared (*20*). The plasmids and primers used for FPA were designed and validated according to a previously published study (*20*). RNA transcription templates for all constructs were generated using a commercial T7 RNA-polymerase forward primer (Eurofins Genomics). All constructs are listed in table S1.

### mRNA transcription and purification

DNA templates were transcribed *in vitro* in a transcription buffer containing 40 mM Tris-HCl (pH 7.5), 15 mM MgCl2, 2 mM spermidine, 10 mM NaCl supplemented with GMP (5 mM), NTPs (3 mM), DTT (10 mM), 1/10 volumes of PCR products, pyrophosphatase (0.005 U/µl), T7 polymerase (0.8%) and RNase inhibitor (0.2 U/µl). Transcription was carried out at 37 °C for 3 h. Following transcription, mRNA was purified using anion-exchange chromatography on a HiTrap Q HP 5 ml column (GE Healthcare). mRNA was eluted using a linear NaCl gradient (0.3–1.5 M NaCl) over 20 column volumes. mRNA-containing fractions were collected and precipitated by adding 1/10 volumes of 20% potassium acetate (pH 5.0) and 2.2 volumes of ethanol at −20°C. The mRNA was pelleted by centrifugation at 4000 rpm for 1 h at 4°C and resuspended in RNase- and DNase-free water. The yield and integrity of the mRNA was assessed using 10% polyacrylamide gel electrophoresis with 8M urea and the absorbance at 260 nm.

### *In vitro* translation and ribosome-nascent chain (RNC) purification

Translation was carried out according to established protocols (*43*) with the following modifications. 70S initiation complex (IC) was assembled by incubating 70S ribosomes (0.5 µM) with a 1.5-fold excess of IF1, IF2, IF3 (0.75 µM each), 1.5-fold excess of BodipyFL-[^3^H]Met-tRNA^fMet^ (0.75 µM) or Atto655-[^3^H]Met-tRNA^fMet^ (0.75 µM), and 4-fold excess of mRNA (2 µM) in HAKM7 buffer and incubating at 37 °C for 45 min. Bpy-[^3^H]Met-tRNA^fMet^ was used to establish efficient *in vitro* translation, stopped-flow experiment, and the limited proteolysis, Atto655-[^3^H] Met-tRNA^fMet^ was used for PET-FCS measurements.

The ternary complex (TC) was prepared by incubating EF-G (4 µM), EF-Ts (0.2 µM), EF-Tu (25 µM), and aa-tRNA (300 µM), DTT (2 mM), GTP (2 mM), phosphoenolpyruvate (PEP) (6 mM), and pyruvate kinase (PK) (0.5 mg/mL) in HAKM7 buffer. All components except aa-tRNA were pre-incubated at 37 °C for 15 min, followed by addition of aa-tRNA and incubation at 37°C for 1 min to form the EF-Tu–GTP–aa-tRNA complex (TC). Translation reactions were performed in HiFi buffer by combining TC (300 µM) and IC (0.04 µM) and incubating at 37°C for 5 min. For truncated mRNAs lacking the stop codon, the translated nascent chains remained bound to ribosomes, forming RNCs.

Translation efficiency was analyzed using Tris-tricine SDS PAGE. Translation reactions containing Bodipy-FL-labeled nascent chains were quenched by adding 1/5 volume of 2 M NaOH, followed by incubation at 37°C for 30 min to release the nascent peptides. The reaction was neutralized by adding an equal volume of 2 M HEPES. For Atto655-containing translation products, 1/10 volume of hydroxylamine (1.5 M) was added instead of NaOH, followed by incubation at 37°C for 1 h. SDS sample buffer (50 mM Tris-HCl pH 6.8, 4% w/v SDS, 2% v/v 2-mercaptoethanol, and 12% w/v glycerol) was added at a 1:1 ratio. Samples were heated at 70°C for 10 min, and peptide products were separated on Tris-Tricine gel. Fluorescent peptides were visualized using an Amersham™ Typhoon™ RGB fluorescence scanner (Cytiva) with excitation wavelengths of 488 nm for Bodipy FL and 680 nm for ATTO-655.

For PET FCS experiments, RNCs were purified by ultracentrifugation through a 1.1 M sucrose cushion in HiFi buffer using a TLA-100 rotor (Beckman Coulter) at 68,000 rpm for 1 h at 4°C. The RNC pellet was resuspended in HiFi buffer, and RNC concentration was determined by liquid scintillation counting of the ^3^H-label using a Perkin Elmer, Tri-Carb 3110 TR Scintillation Liquid Analyzer. Purified RNCs were flash-frozen in liquid nitrogen and stored at −80°C.

### PET-FCS

PET-FCS measurements were performed on the MicroTime 200 system (PicoQuant) built on a modified Olympus IX 73 confocal microscope equipped with 60x water immersion objective Olympus UPlanSApo 1.2 N.A. (Olympus UPlanSApo). A excitation 636.5 nm laser (operated in continuous wave mode) was adjusted to ∼ 40 µW to prevent photobleaching and decrease the formation of Atto655 triplet state (*20*). Emitted fluorescence signal was focused through a pinhole, split 50/50 by a beam splitter, passed through a 690/70 nm band pass filter and detected by two single-photon avalanche photodiodes (SPAD). Cross-correlation of fluorescence trace was used to eliminate SPAD after-pulsing effects.

To ensure single molecule detection within the confocal detection volume (∼1 femtoliter), purified RNCs were diluted to ∼10 nM in HiFi buffer. Each measurement used 50 µl of the RNC sample (∼10 nM). Where indicated, TF (3 µM) was added to the RNC sample. All experiments were conducted at least 2-3 times with RNCs from independent preparations. For each RNC solution, four replicate measurements were recorded at consecutive 10-min intervals at room temperature (22°C). Fluorescence traces were analyzed by the SymPhoTime 64 software (PicoQuant). The autocorrelation function (ACF) was calculated using 314 time points between 0 and 1 s. The reliability of each measurement was confirmed by comparing replicates, with overlapping diffusion curves indicating RNC stability during measurement. As the lateral dimension of the detection volume is significantly larger than the axial dimension, a 2D single-species diffusion model was used to fit the ACF:

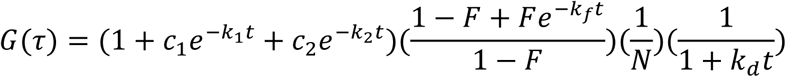

where *k*1 and *k*2 represent relaxation rate constants, and *c*1 and *c*2 their respective amplitudes. *kf* denotes the rate constant of triplet state decay, with F as the corresponding amplitude. *kd* is the diffusion coefficient and *N* represents the average number of molecules within the focal volume. Across all constructs, RNC diffusion and triplet state decay rates are similar, with relaxation times of ∼1 ms and ∼40 µs, respectively. The fitted parameter *N* was used to normalize the ACF to *N*=1.

After refitting the ACF, the Y-intercept *G*(0) used to quantify the proportion of fluorescent molecules:

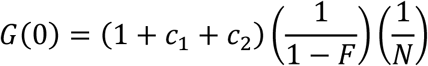

The dynamicity of nascent chains (*DNC*) was calculated as:

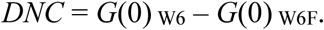

The resulting dimensionless value provides a qualitative estimate for the fraction of dynamic nascent chains. The effect of mutations was estimated as:

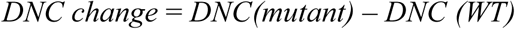

A detailed analysis of the fluctuation dynamics from the rate constants of the ACF was not performed, because the exact values of the calculated elemental rate constants are somewhat model-dependent and therefore do not provide additional insights beyond the qualitative comparison of WT and mutants based on the *G*(0) values.

### Limited proteolysis

All measurements were performed using RNCs labeled with N-terminal Bodipy FL. Proteolysis was initiated by mixing RNCs (0.02 µM) and thermolysin (0.5 ng/µl, Promega) in HiFi buffer supplemented with CaCl2 (0.25 mM). Reactions were incubated for the indicated times at 37°C and quenched by adding 1.75 volumes of quenching solution containing 85 mM Tris-HCl (pH 6.8), 6% SDS, 1.7% 2-mercaptoethanol, 17% glycerol and 0.3 M NaOH, followed by incubation at 85°C for 15 min (*9*). To prevent significant fluorophore bleaching pH was adjusted to neutral by adding 2M HEPES. Translation products were analyzed by Tris-Tricine SDS-PAGE and visualized via Bodipy fluorescence using the Amersham™ Typhoon™ RGB scanner (Cytiva). Band intensities on SDS-PAGE were quantified using the Multi Gauge software, and digestion time courses were evaluated by single-exponential fitting using Graphpad Prism (Version 9.0.0).

### Data analysis of the force profile assay (FPA)

FPA mRNA constructs contained the N-terminal HemK codon sequence of varying length in one or two-codon increments, followed by 17 codons of the SecM arrest peptide, and 20 codons of protein CspA (table S1). Under low-force conditions, translation is arrested at the SecM sequence, resulting in the formation of an arrested translation product (AR) (*20*). However, if nascent peptide folding generates force, translation arrest is alleviated, leading to the production of a longer full-length peptide (FL) (fig. S6). Translation products were separated by SDS-PAGE, and the fraction of the FL product was calculated as:

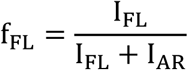

where IFL and IAR represent the band intensity of the respective products. Experiments were repeated three times, and mean ± SD values were calculated.

### Stopped-flow experiments

To monitor PET between the N-terminal Bodipy-FL and Trp residues in the polypeptide chain during real time translation, stopped-flow PET experiments were performed at 37°C using an SX-20MV device (Applied Photophysics). Excitations was set to 470 nm (slits width = 2 mm, bandpass of 4.65 nm/mm), and emission was detected through a KV500 cut-off filter (Schott). Equal volumes of purified IC and freshly prepared TC were rapidly mixed to initiate the translation reaction, and fluorescence changes were recorded with 4000 time points over 2 min on a logarithmic scale. Fluorescence was normalized to the initial fluorescence, calculated from the averaged signal between 0.001 and 0.005 s. For each experiment, at least two biological replicates with six technical replicates each were averaged. PET efficiency was calculated according to the equation: *E*PET = 1 – FW6/ FW6F, where FW6 and FW6F are fluorescence of the W6 and W6F constructs, respectively.

**Fig. S1.**
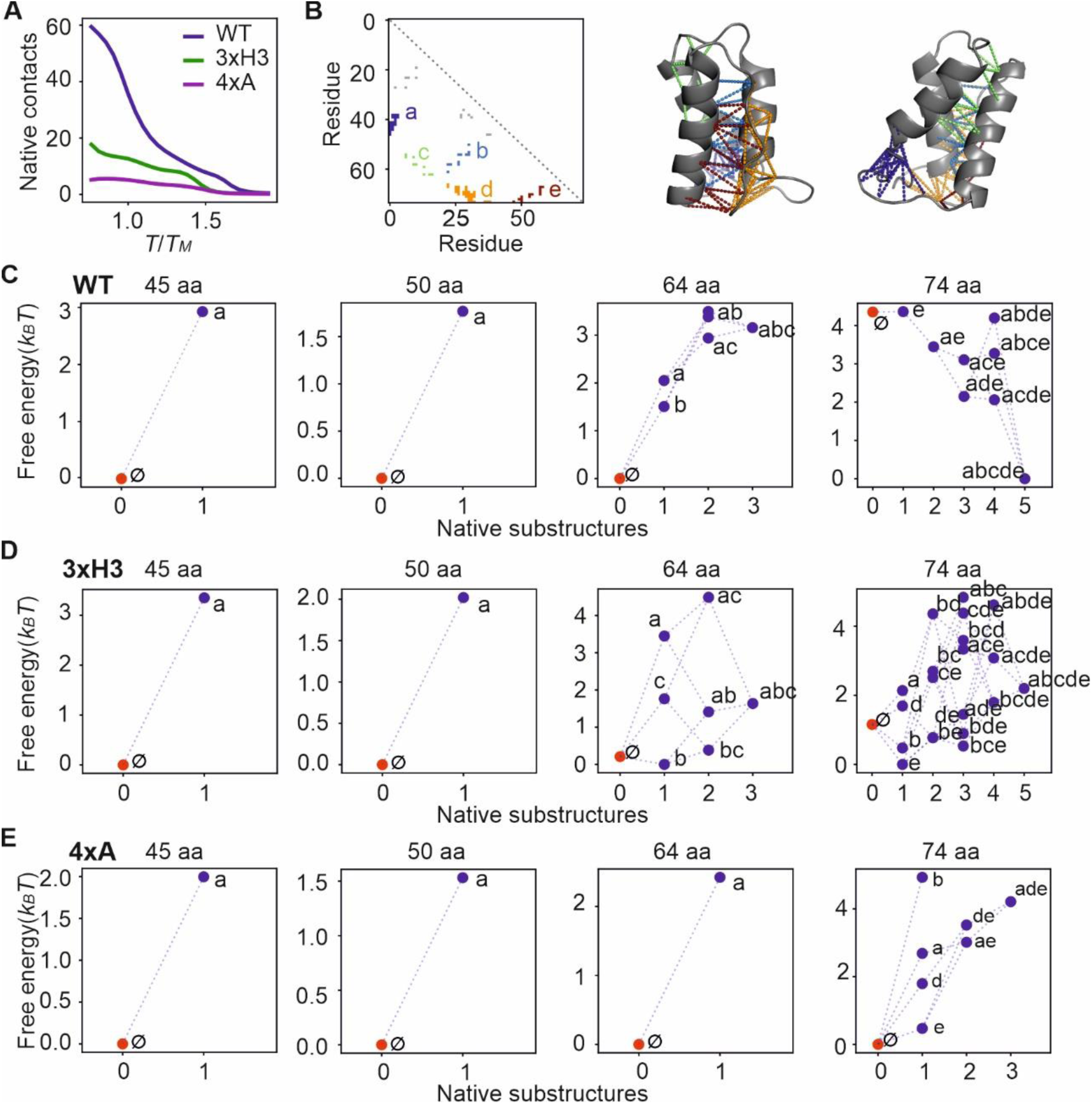
Simulation of equilibrium free energy landscapes during synthesis reveals the transition from non-native to native-like structure. (**A**) Simulated thermal melting curves showing the equilibrium fraction of native contacts as a function of temperature for full-length (74 aa) WT HemK NTD and the 3xH3 and 4xA mutants. Temperature is normalized to the experimentally determined melting temperature (Tₘ) of WT HemK NTD (323 K). Simulation results are presented at T = 0.86 Tₘ, corresponding to in vitro experimental conditions at 293 K. (**B**) Contact map of full-length HemK (74 aa), with folding substructures identified as groups of adjacent native contacts that cooperatively form and break. Substructures are color-coded and labeled alphabetically. Gray-shaded points indicate isolated contacts that do not belong to any substructure and are expected to form and break non-cooperatively. To the right, two views of the equilibrated HemK structure are shown, with contacts corresponding to substructures marked as colored dashes matching the contact map. (**C**) Free energy landscapes of WT HemK as a function of topological configuration, defined by the subset of formed substructures, at different protein lengths. Each dot represents a single configuration, positioned on the X-axis by the number of substructures formed and on the Y-axis by its free energy (in kBT). Red dots indicate configurations with no native structures (∅). Dots are connected if they differ by a single substructure, representing potential folding pathways. Notably, these configurations represent only native folding; for example, a snapshot containing a substantial number of non-native contacts is classified within a configuration with few or no substructures (∅ or a). At intermediate lengths (left panels), the landscape is dominated by non-native states, whereas at full length (right panel), native folding becomes thermodynamically favorable. (**D, E**) Free energy landscapes for the 3xH3 (D) and 4xA (E) mutants. Native-like folding is unfavorable at all lengths. However, in the 3xH3 mutant, native-like intermediates (e.g., bce) are more favorable at full length compared to the 4xA mutant. In the 4xA mutant, the free energy difference between the native state (abcde) and the most stable non-native state (∅) exceeds the 5 kBT cutoff and is therefore not shown. Although 3xH3 residues are solvent-exposed in the native structure, they destabilize the full-length native state (D). In our simulations, local structural fluctuations transiently bring these hydrophobic residues into the core, stabilizing the native state in WT, whereas mutations disrupt this effect.

**Fig. S2.**
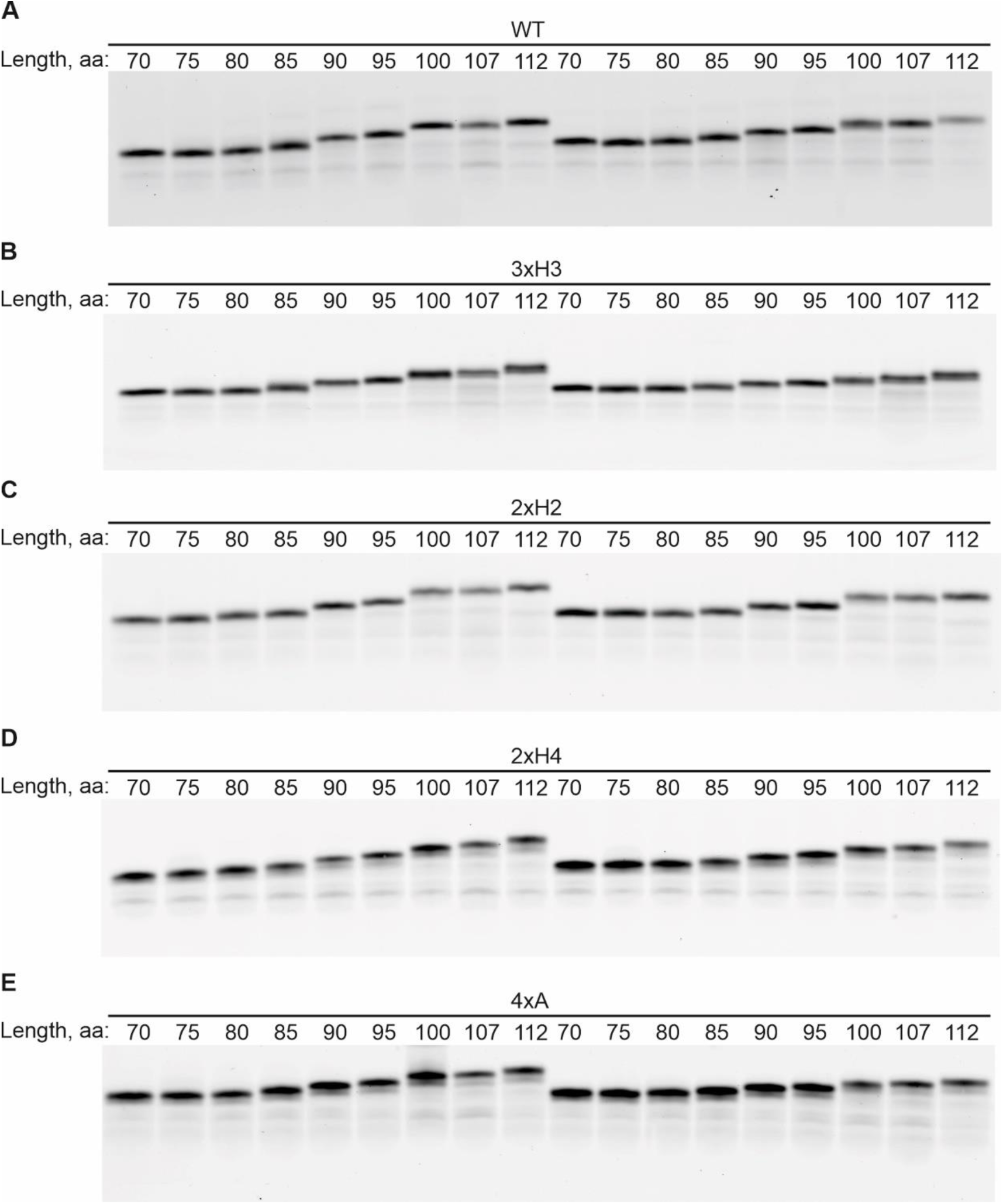
*In vitro* translation efficiency of WT, 3xH3, 2xH2, 2xH4, and 4xA mRNA constructs of different lengths. Translation products were separated using Tris-Tricine SDS-PAGE and visualized via the fluorescence of the BODIPY-FL reporter attached to the N-terminus of the peptide. The aa lengths of the HemK constructs are indicated. (**A**) WT W6 and WT W6F. (**B**) 3xH3 W6 and 3xH3 W6F. (**C**) 2xH2 W6 and 2xH2 W6F. (**D**) 2xH4 W6 and 2xH4 W6F. (**E**) 4xA W6 and 4xA W6F.

**Fig. S3.**
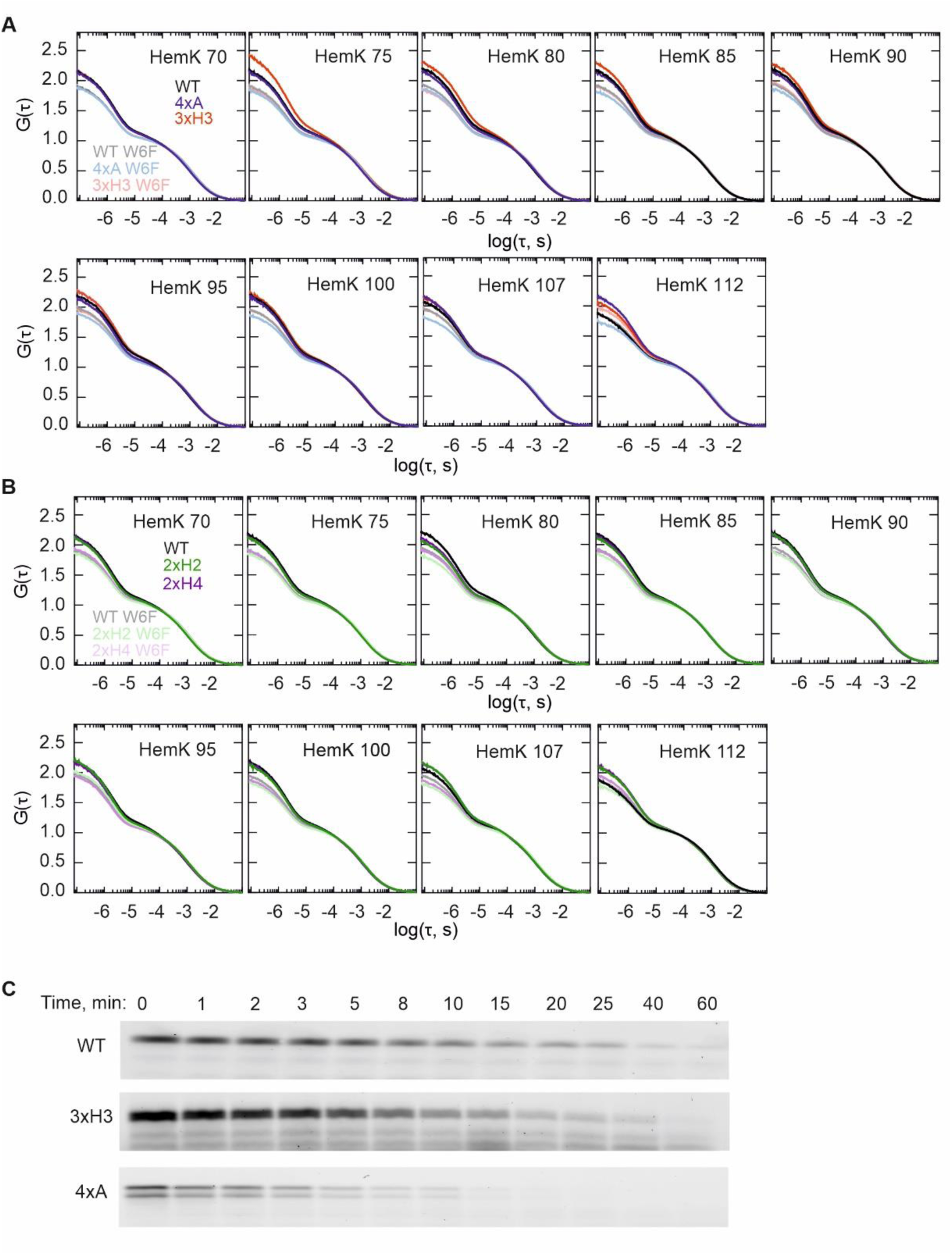
PET-FCS autocorrelation and proteolysis. (**A, B**) PET-FCS autocorrelation function (ACF) curves for WT, 3xH3, and 4xA (**A**), as well as 2xH2 and 2xH4 (**B**), nascent chains at the indicated chain lengths. ACFs were obtained from at least two biological replicates with a minimum of four technical replicates each (N=8). (**C**) Time courses of thermolysin proteolysis for WT, 3xH3, and 4xA 112-aa nascent chains, highlighting differences in protease sensitivity.

**Fig. S4.**
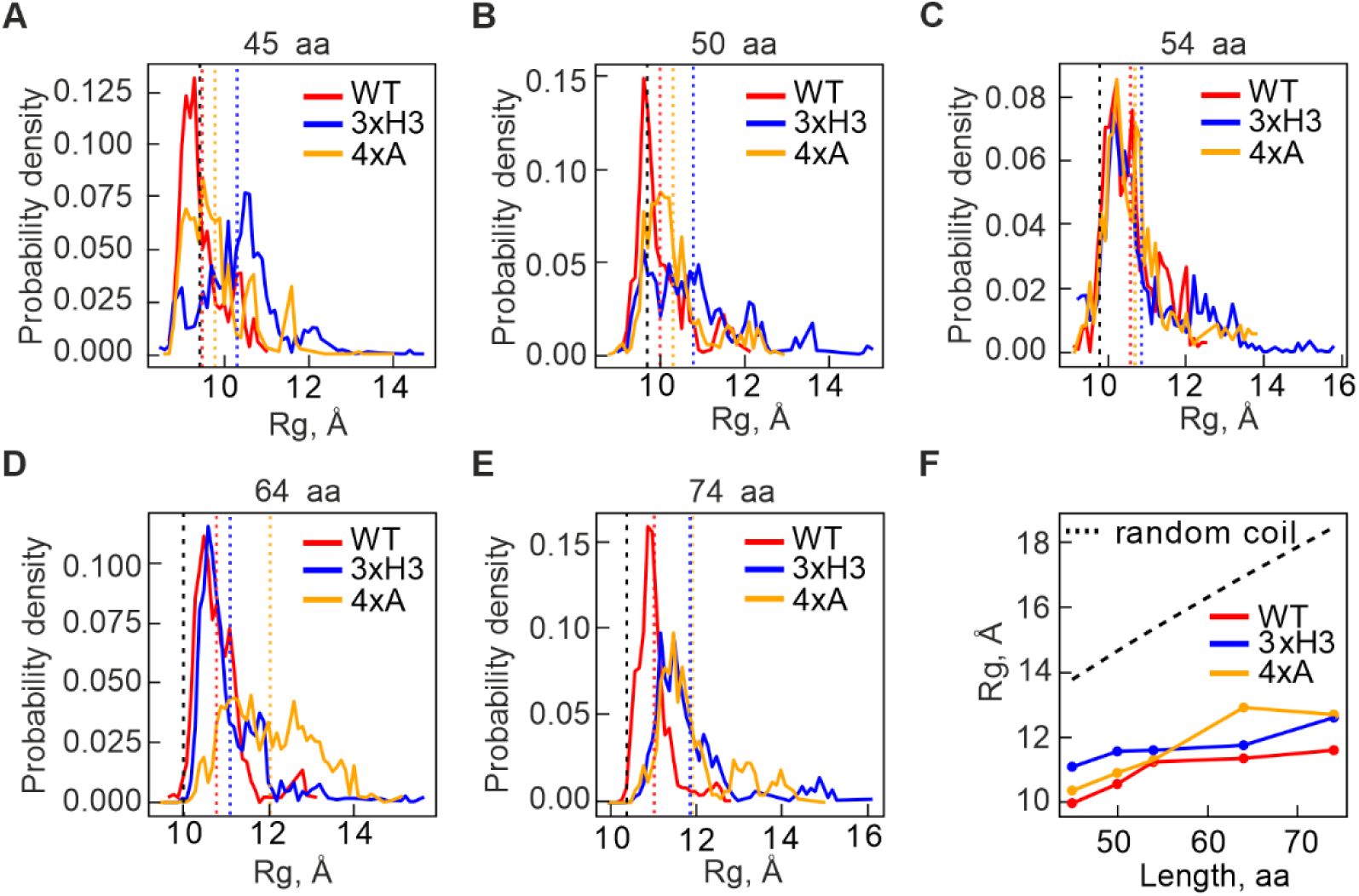
Non-native interactions maintain compactness of early folding intermediates. (**A**-**E**) Probability density plots of the radius of gyration (Rg) as a function of peptide length for WT HemK (red), 3xH3 mutant (blue), and 4xA mutant (orange). Vertical dashed lines indicate the average Rg for each construct in the corresponding color, while the expected Rg for native-like folding at each length is shown as a vertical dashed black line. Peaks in the distributions suggest the presence of multiple interconverting states with distinct average Rg values, leading to non-symmetric Rg distributions rather than the symmetric distributions expected for a single folding state. (**F**) Average Rg as a function of chain length for WT, 3xH3, and 4xA mutants, as well as expected value for a random coil control, illustrating differences in compactness due to non-native interactions in the mutants.

**Fig. S5.**
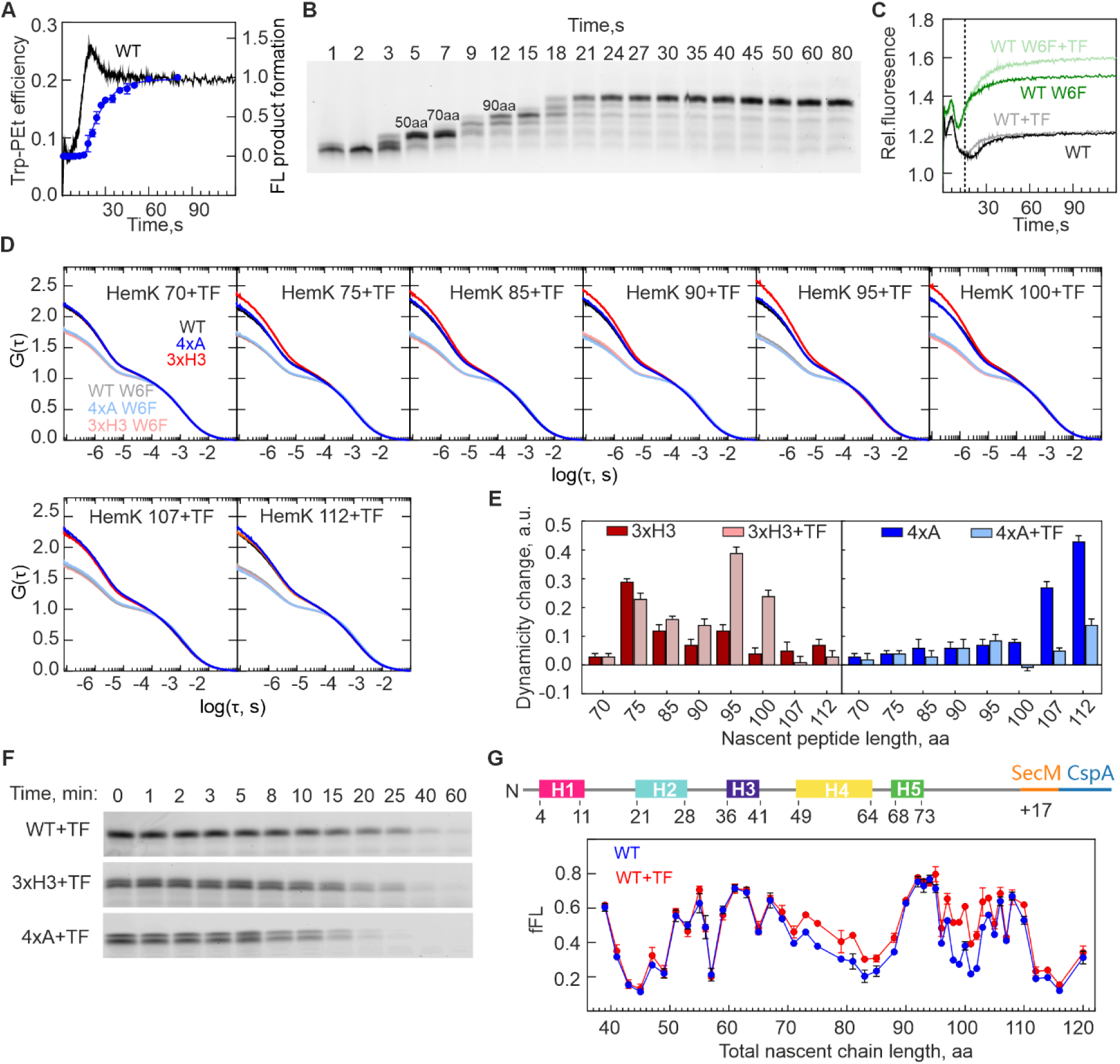
Cotranslational folding during real time translation and the effect of TF. (**A**) Stopped-flow time course of HemK NTD cotranslational compaction, monitored by PET between BodipyFL-Met1 and Trp6 in the nascent chain (black), together with the time course of full length product formation (blue). (**B**) Translation time course for WT mRNA. The lengths of transiently-accumulating translation are indicated. (**C**) Changes in BOF-Met fluorescence during translation of WT mRNA (black) and W6F mRNA (green) in the absence of TF, and in the presence of TF (gray and light green, respectively). The dashed line marks the time point of TF’s initial interaction with the nascent peptide. (**D**) PET-FCS ACF curves at the indicated nascent peptide length in the presence of TF. (**E**) DNC changes relative to WT for 3xH3 (red) and 4xA (blue) mutants in the absence and presence of TF. Error bars represent SEM. (**F**) Time courses of thermolysin proteolysis for WT, 3xH3, and 4xA 112-aa nascent chains in the presence of TF, highlighting differences in protease sensitivity. (**G**) FPA profile of the WT nascent peptide in the presence (red) and absence (blue) of TF. The total nascent chain length includes both the HemK nascent peptide and the SecM stalling sequence. Data represent the mean ± SD from three biological replicates. A schematic of the FPA experiment construct is shown on top (see Methods for details).

**Fig. S6.**
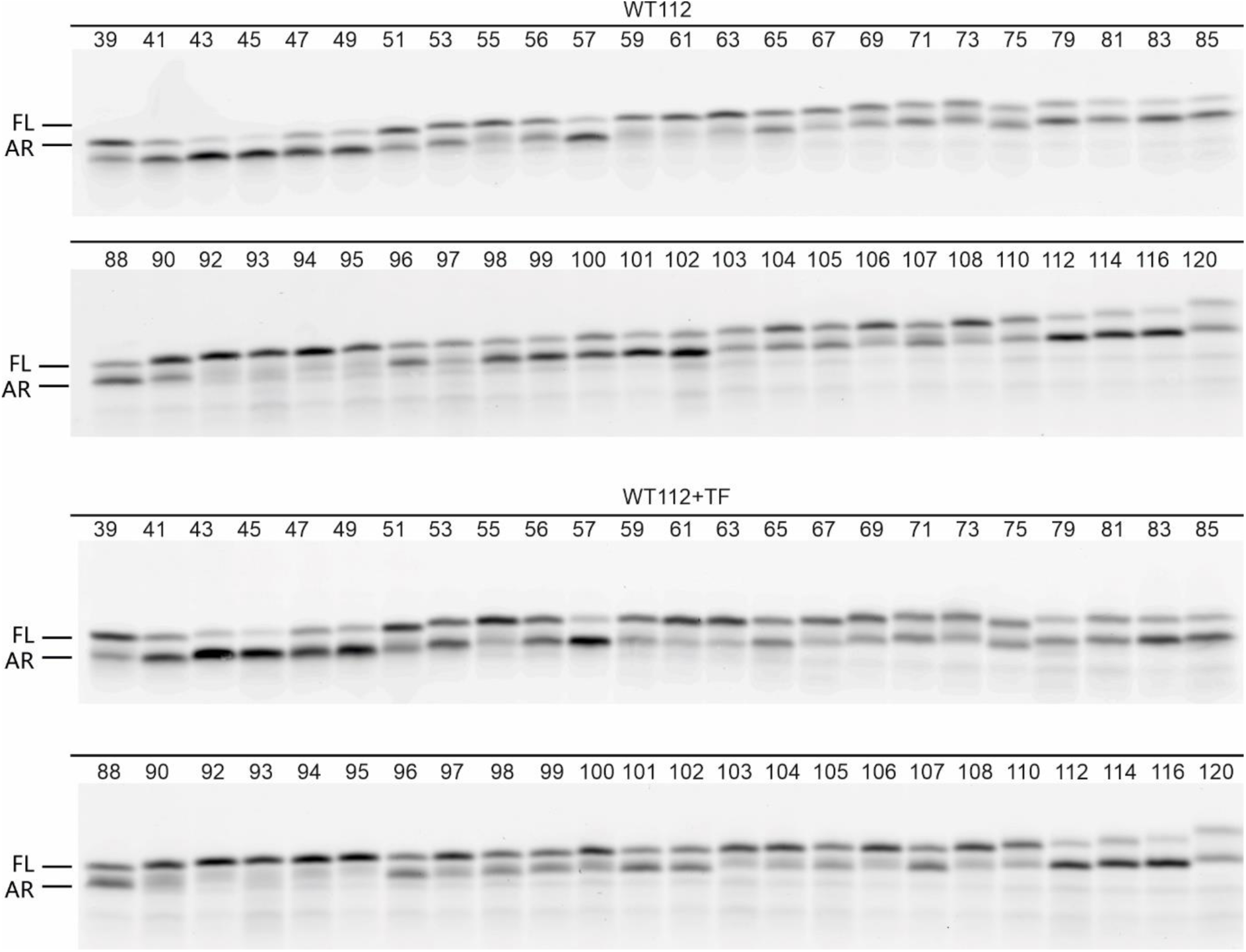
FPA analysis. SDS-PAGE of translation products shows two bands which were used to calculate the fraction of full length (FL) compared to total FL and arrest (AR) peptide in Fig 4E.

**Table S1.**
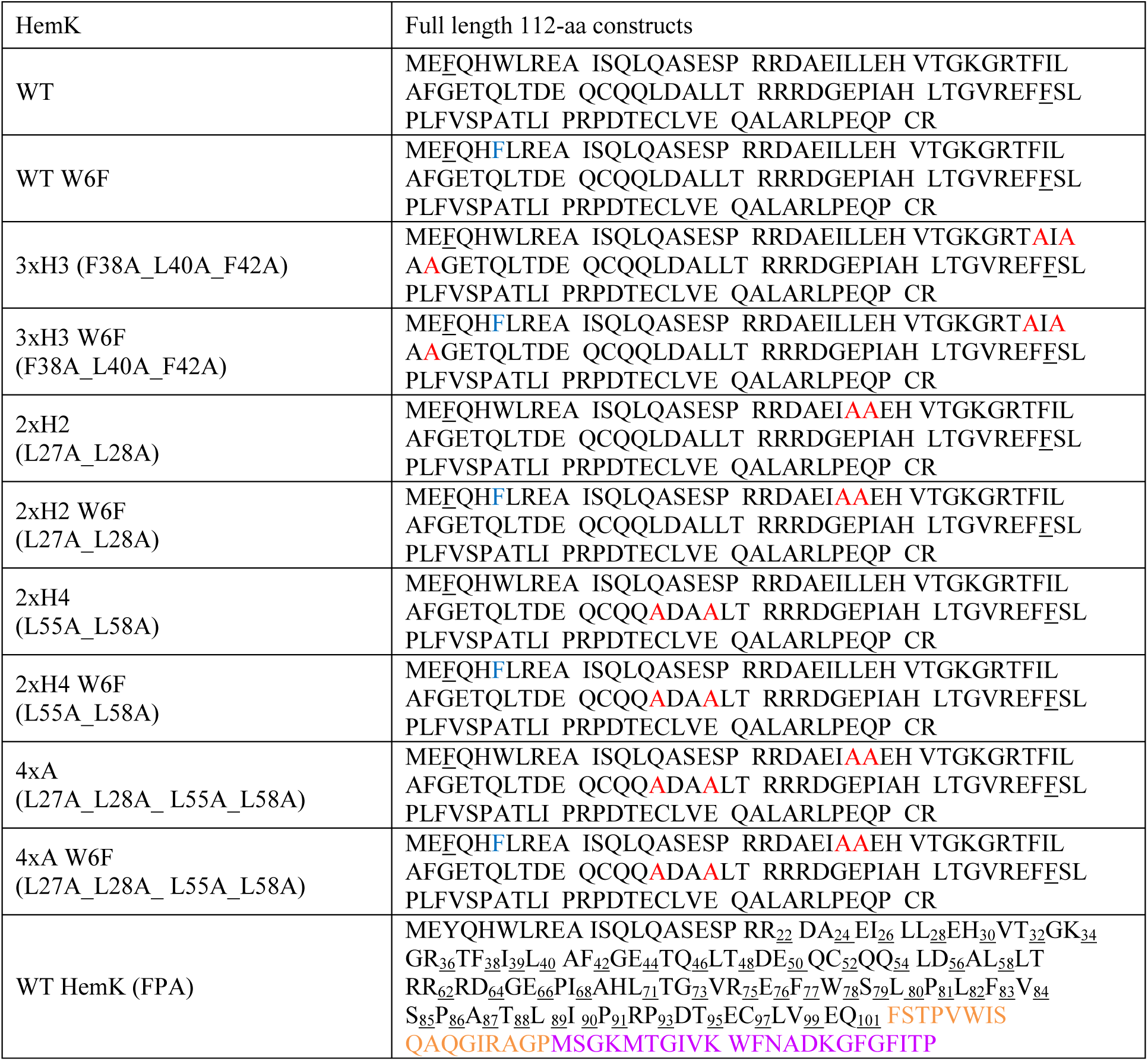
List of aa sequences of protein constructs studied by PET-FCS and FPA.

**Table S2.**
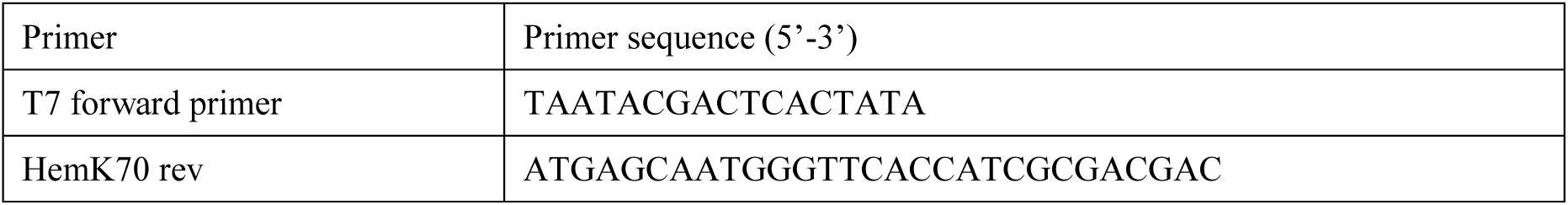
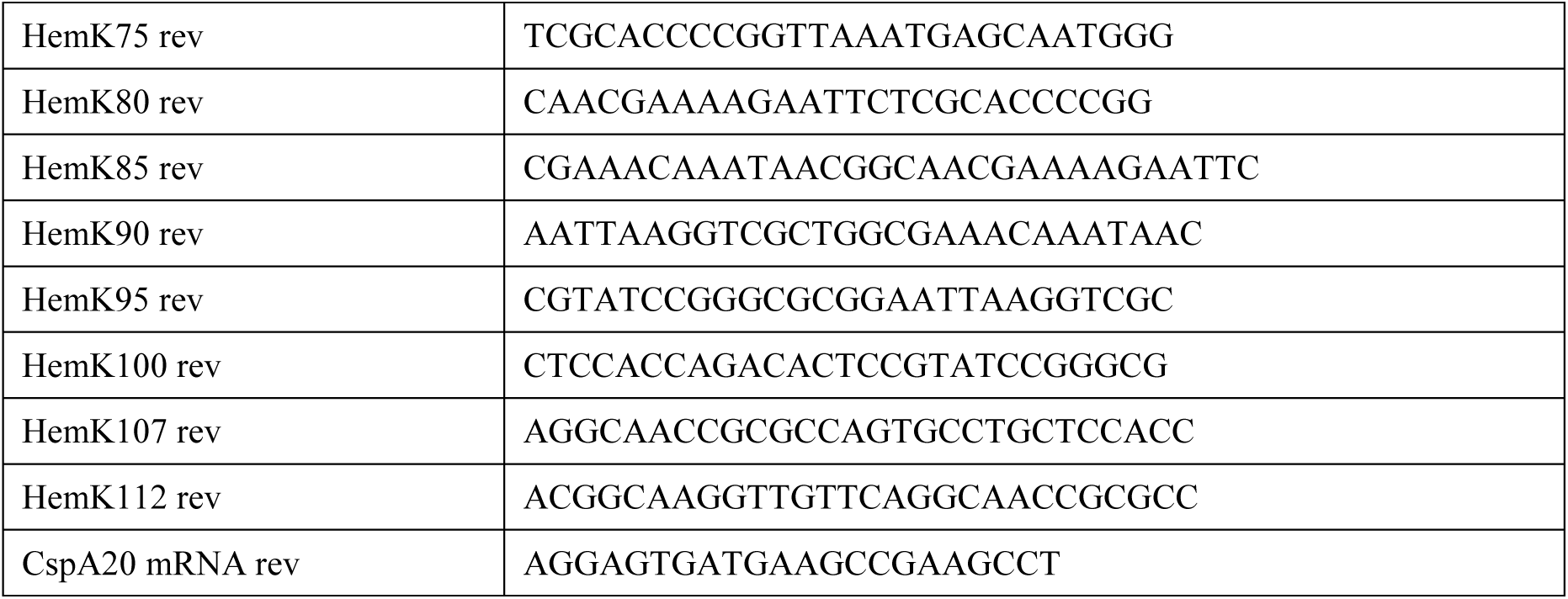
Primers used to generate mRNA transcription templates.

